# Diversity of Short Linear Interaction Motifs in SARS-CoV-2 Nucleocapsid Protein

**DOI:** 10.1101/2023.08.01.551467

**Authors:** Peter Schuck, Huaying Zhao

## Abstract

Molecular mimicry of short linear interaction motifs has emerged as a key mechanism for viral proteins binding host domains and hijacking host cell processes. Here, we examine the role of RNA-virus sequence diversity in the dynamics of the virus-host interface, by analyzing the uniquely vast sequence record of viable SARS-CoV-2 species with focus on the multi-functional nucleocapsid protein. We observe the abundant presentation of motifs encoding several essential host protein interactions, alongside a majority of possibly non-functional and randomly occurring motif sequences absent in subsets of viable virus species. A large number of motifs emerge *ex nihilo* through transient mutations relative to the ancestral consensus sequence. The observed mutational landscape implies an accessible motif space that spans at least 25% of known eukaryotic motifs. This reveals motif mimicry as a highly dynamic process with the capacity to broadly explore host motifs, allowing the virus to rapidly evolve the virus-host interface.

## INTRODUCTION

Short linear interaction motifs (SLiMs) are stretches of several amino acids that serve as microdomains in intrinsically disordered regions (IDRs) mediating weak, transient protein-protein interactions with target domains (Sangster et al., 2022). SLiMs have emerged as a ubiquitous modules in the organization of protein-protein interaction networks. For example, SLiMs designate substrates for post-translational modification including kinases and phosphatases, target cellular localization, and control docking to adaptors, assembly of signaling complexes, and recruitment of enzymes to multi-protein complexes (Manna et al., 2018; Mohapatra and Dixit, 2022; Peti and Page, 2013; Ragusa et al., 2010; Shi et al., 2023; Sorgeloos et al., 2022; Srivastava et al., 2023; Wang et al., 2016). The number of eukaryotic SLiM classes is in the hundreds and rapidly expanding, and it has been estimated there may be more than a hundred thousand of instances of such motifs in the human proteome (Davey et al., 2023; Kumar et al., 2022; Tompa et al., 2014).

Molecular mimicry of SLiMs is a key mechanism for viral proteins to hijack and modulate host cell processes and is broadly exploited among many viruses (Davey et al., 2011; Hagai et al., 2014; Mihalič et al., 2023). Therefore, the distribution and evolution of viral motifs is of significant interest in the search for broad-spectrum anti-viral drug targets (Shuler and Hagai, 2022; Simonetti et al., 2023). It has been proposed that the evolution of SLiMs is facilitated by their compact size, in combination with the unusually high abundance of IDRs in viral proteins; the latter, by virtue of fewer constraints, generally exhibiting high mutation frequencies that would allow the efficient *de novo* formation of SLiMs through as little as a single amino acid change (Brown et al., 2002; Davey et al., 2015; Fuxreiter et al., 2007; Gitlin et al., 2014; Neduva and Russell, 2005; Tokuriki et al., 2009). The conservation of disorder has been hypothesized to facilitate change of interaction partners and thereby provide an evolutionary advantage (Mosca et al., 2012). In eukaryotes, the generation and loss of SLiMs has been confirmed in sequence analyses of evolutionarily related species, as well as in rare random mutations individual human patients (Cordeddu et al., 2009; Davey et al., 2015). In viruses, substantial heterogeneity of motif content has been observed across different viral families, with instances of convergent evolution, pointing to a highly dynamic repertoire of motif usage and evolutionary adaptation (Hagai et al., 2014).

On the other hand, a salient feature of RNA-viruses is their high intracellular and intrahost sequence diversity due to their evolved low transcription fidelity and resulting quasispecies nature (Domingo, 2019; Eigen, 1993; Holland and Domingo, 1997; Lauring and Andino, 2010). How this sequence diversity impacts the virus-host interface and the evolution of SLiMs has remained unexplored. An opportunity to study this question recently arose with the unprecedented, vast collection SARS-CoV-2 genomes in GISAID (Elbe and Buckland-Merrett, 2017). While it provides a basis for monitoring the evolution of mutations distinguishing emergent clades, the repository contains a majority of random transient mutations that exhaustively explore the mutational landscape of viral proteins (Bloom et al., 2023; Saldivar-Espinoza et al., 2023; Zhao et al., 2022). In the present work, we exploit the diversity of SARS-CoV-2 sequences and examine the distribution of viral motif content on the infected host population level, which we propose may serve as a model also for intrahost viral motif diversity.

Eukaryotic motif mimicry for cell entry has been found in the RBD of the spike protein of SARS-CoV-2 (Mészáros et al., 2021). In the present work we focus on the nucleocapsid (N-)protein, which has several favorable properties as platform for SLiMs, having the highest expression level of all SARS-CoV-2 proteins at ≈1 % of total protein in infected cells (Tugaeva et al., 2021), and containing three IDRs spanning nearly half of the protein (**Figure 1A**). Even though it provides a major antigen, it is not immuno-dominant as the spike protein. Besides its eponymous structural role in viral assembly (Carlson et al., 2022; Masters, 2019; Zhao et al., 2023), N-protein is highly multi-functional with a large host interactome (Gordon et al., 2020; Kruse et al., 2021; Wu et al., 2023; Zheng et al., 2021), including interactions with proteins of the type 1 interferon signaling pathway (Li et al., 2020; Mu et al., 2020; Yelemali et al., 2022), the inflammasome (Pan et al., 2021), complement activation (Gao et al., 2022), lipid metabolism (Yuan et al., 2021), and expression and binding to cytokines (Karwaciak et al., 2021; López-Muñoz et al., 2022). Among interactions described in greatest biophysical detail are the complex formation with G3BP1 leading to rewiring of stress granules (Biswal et al., 2022; Kruse et al., 2021; Yang et al., 2023), binding of 14-3-3 (Eisenreichova and Boura, 2022; Tugaeva et al., 2021, 2023), and the interaction with host kinases leading to extensive phosphorylation particularly in the linker IDR of intracellular N-protein (Carlson et al., 2020; Syed et al., 2023; Yaron et al., 2022). Other posttranslational modifications include ubiquination, proteolytic cleavage, sumoylation, and ADP-ribosylation (Fung and Liu, 2018; Madahar et al., 2023; Mao et al., 2023).

**Figure 1.**
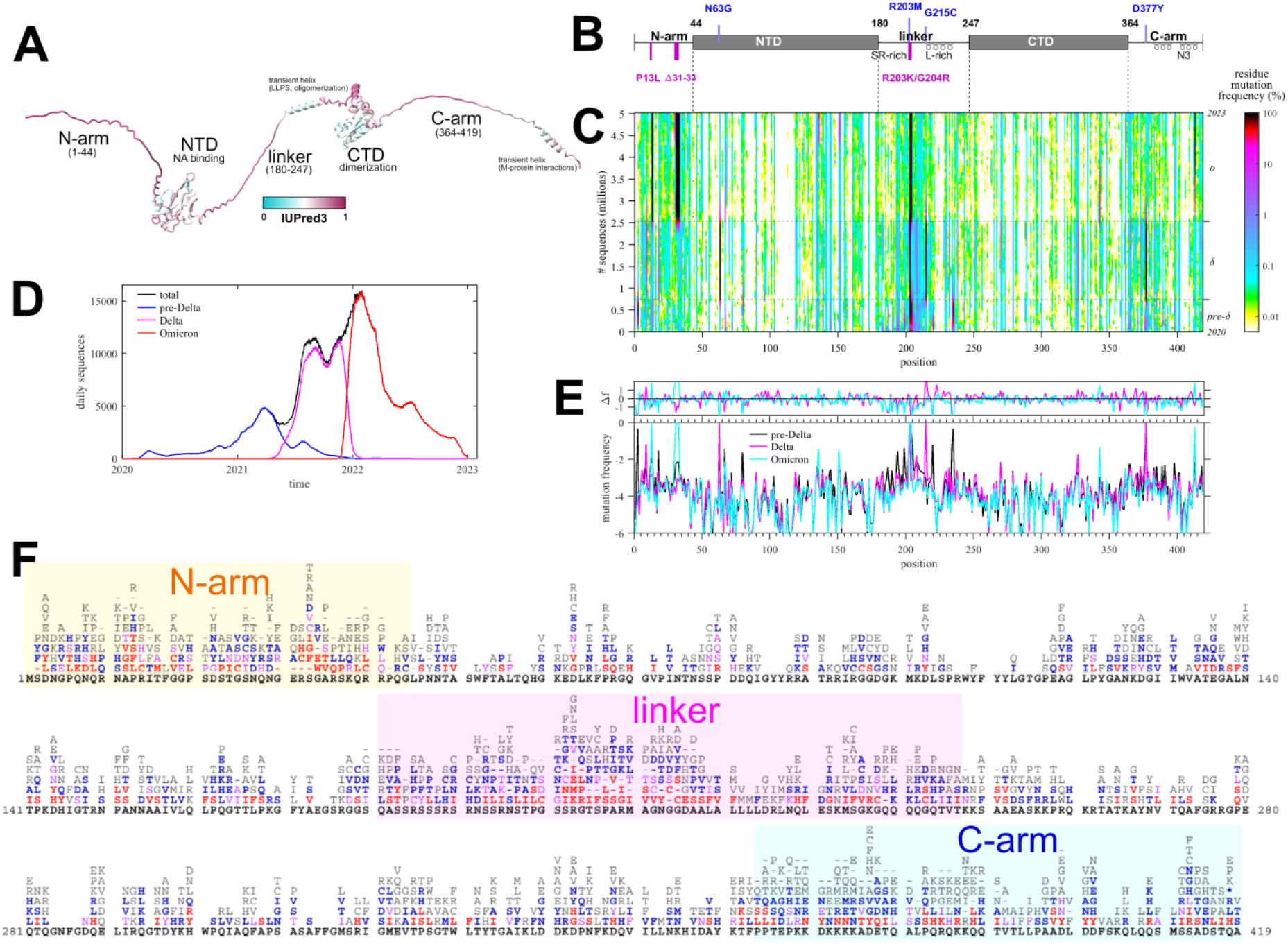
Mutational landscape of N-protein. (A) Predicted AlphaFold structure of N-protein, with extended IDRs for clarity, colored according to the disorder score. Labeled are the folded domains (NTD, CTD) and the three IDRs (N-arm, linker, and C-arm) with known assembly functions. (B) Schematic organization of N-protein with defining mutations of Delta (clade 21J, blue) and Omicron variant (magenta). (C) Time-dependent mutation frequencies (colors) at different positions (abscissa) plotted as a function of cumulative sequences (ordinate). The dotted horizontal lines are the time-points where Delta- and Omicron-variants rose to majority, as indicated in the history of daily deposited sequence numbers grouped by pre-Delta (comprising ancestral, Alpha, Beta, and other variants predating Delta 21J), Delta, and Omicron variants (D). (E) Average mutation frequencies of residues along N-protein positions, grouped by major variants as in (D). The upper panel shows the difference in average mutation frequencies of Delta and Omicron relative to the pre-Delta group. (F) Detailed list of observed amino acid mutations in each position, colored by the number of observed instances (≥ 1,000 red, ≥ 500 magenta, ≥ 100 blue, ≥ 10 grey, ancestral black).

The basis for the present work is a dataset of ≈5 million SARS-CoV-2 consensus sequences, which we first characterize regarding the mutation frequencies and mutational landscape of N-protein over time. We have recently exploited the exhaustive map of viable amino acid mutations as a tool in the structural analysis of assembly roles of the linker IDR (Zhao et al., 2022, 2023). For the analysis of motif content we examine the ancestral consensus sequence and utilize a previously introduced statistical approach (Hagai et al., 2014) to identify possibly random motif instances. We then determine the distribution of motif content in the ensemble of viral sequences sampled across the infected host population. This analysis reveals a highly dynamic motif presentation where most of the ancestral motifs are abandoned in at least some sequence subsets, while in others large numbers of new motifs arise that potentially interact with various host pathways. Finally, based on the mutational landscape, we estimate a measure of the available sequence space and corresponding accessible motif space, which suggests that even for the smallest N-protein IDR a large fraction of known motifs may be presented. While many or most of these may not be functionally interacting with host proteins, it reveals a mechanism for extensive probing of viral proteins for potentially beneficial host protein interactions. We discuss possible implications for the virus-host interface considering intrahost and intracellular sequence diversity.

## RESULTS

### The mutational landscape of SARS-CoV-2 nucleocapsid protein

The present study is based on 5.06 million high-quality consensus SARS-CoV-2 sequences retrieved on January 20, 2023 from Nextstrain (Hadfield et al., 2018). Due to their origin from COVID19 patient samples we may assume these to be sequences of viable and infectious virus. The sequences exhibit significant diversity, with ≈56% of sequences being distinct, ≈30% unique, and ≈8% spatio-temporally distant repeats. The set contains ≈43 million instances of N-protein mutations relative to the ancestral Wuhan-Hu-1 isolate. Most of the mutations are different from the defining substitutions of the variants of concern (**Figure 1B**), and instead occur transiently and are distributed across ≈92% of all N-protein residues. This suggests these mutations may have a significant impact on the N-protein motif repertoire.

**Figure 1C** shows the frequency of these evolutionarily inconsequential mutations at different positions and as a function of time (where time is plotted in the ordinate transformed to a cumulative sequence number to compensate for uneven sampling frequency (**Figure 1D**)). As may be discerned from the barcode-like vertical patterns in the frequency plot **Figure 1C**, the local mutation frequencies are in first approximation constant outside the defining variant substitutions, with minor evidence of limited stochastic transmission events causing temporal variation. By contrast, there is significant structure across different positions: Higher frequencies are generally observed in the IDRs, and lower mutation frequencies correlate with biophysical functional constraints in the folded domains and IDRs (Zhao et al., 2022, 2023). Short of a detailed evolutionary analysis, we may roughly group sequences into three distinct sets of Omicron variants, Delta variants, and those preceding the Delta-variants, which constitute the vast majority of sequences in different periods of time (**Figure 1D**). Their residue mutation frequency pattern is nearly identical outside the defining substitutions, which shows that major constraining biophysical properties of N-protein are unaltered. This is highlighted further in the detailed comparison of average residue mutation frequencies as a function of position among the three groups (**Figure 1E**).

A detailed chart of the observed amino acid mutations at all positions is shown in **Figure 1F**. On average ≈5.5 different amino acids may occupy each position, ranging from zero mutations at 14 of the 37 conserved positions across related coronaviruses, to a maximum of 12 different amino acids that may occupy the most variable positions of the IDRs. Their pattern defines a characteristic mutational landscape, which is similar to that reported previously (Zhao et al., 2022), despite the fact that the current data set includes ≈2.6 million Omicron sequences (separately shown in **Figure S1**) that were not yet available previously. We conclude that the different waves of SARS-CoV-2 variants independently reproduce very similar mutational landscapes, as would be expected for exhaustively sampled N-protein in a steady-state.

The mutational landscape is a comprehensive set of all tolerable, non-lethal mutations (Bloom and Neher, 2023) and as such it reflects detailed biophysical constraints and provides complementary information to traditional structural tools (Zhao et al., 2022, 2023). For example, binding to G3BP1/2 was identified as an essential N-protein function and a crystal structure shows G3BP1 binding the φ-x-F motif in the N-arm (Biswal et al., 2022). Accordingly, F17 is an almost completely conserved residue in the mutational landscape of the otherwise highly variable IDR (**Figure 1F**). Similarly, in the leucine-rich region of the linker IDR the oligomerization of transient helices to form coiled-coils was recently identified as an essential assembly function, and structural requirements for oligomerization were found to be reflected in the nature of the limited set of amino acid mutations in positions 221-233, in the otherwise highly variable linker IDR (Zhao et al., 2023).

Different combinations of N-protein mutations define 40,988 distinct N-protein sequences. Since linear motifs are prevalent in IDRs, we focus on the subset of distinct IDR sequences. Using a threshold condition that each sequence is observed in at least 10 different genomes, there are 512 distinct sequences for N-arm carrying an average of 2.75 mutations, 979 for the linker IDR with an average of 2.90 mutations, and 556 for the C-arm IDR with an average of 1.77 mutations. Each sequence was examined with regard to their SLiM content using the Eukaryotic Linear Motif (ELM) Resource for Functional Sites in Proteins (Kumar et al., 2022), which can search for occurrences of regular expressions of 327 documented motif classes.

### Motif content of the ancestral SARS-CoV-2 N-protein sequence

As a starting point to parse the results we consider first the motif content in the linker IDR of the ancestral Wuhan-Hu-1 sequence. **Figure 2** lists in bold the predicted ancestral motifs and the inset shows their location. Many motif classes occur in multiple instances, their number charted as white crosses. The motif set is dominated by sites for kinases, in particular in the SR-rich region. This is not surprising, considering the high degree of phosphorylation experimentally observed (Yaron et al., 2022). Kinase motifs significantly overlap, which may produce allovalency and allow for cooperativity and increase effective affinity of the sites for kinase binding (Klein et al., 2003). In addition, several other motifs overlap in both the SR-rich and the transiently helical L-rich region of the linker (**Figure 1B**), which may not preclude their function considering the large number of intracellular copies N-protein. A previously reported 14-3-3 motif in the linker (Eisenreichova and Boura, 2022; Tugaeva et al., 2023) is reproduced, and a variety of motifs for different posttranslational modifications and binding functions are found. A similar preponderance of phosphorylation motifs is found in the C-arm (**Figure 3**) and N-arm (**Figure 4**) IDRs (the latter missing the above mentioned G3BP1 binding motif not contained in the ELM database). While some of the other motifs for protein modification and host protein interactions seem plausible, such as those related to de-ubiquination, sumoylation, autophagy, and apoptosis, others appear unlikely to describe real interactions, for example, several glycosylation motifs (Shajahan et al., 2021) and those targeting proteins of different organisms.

**Figure 2.**
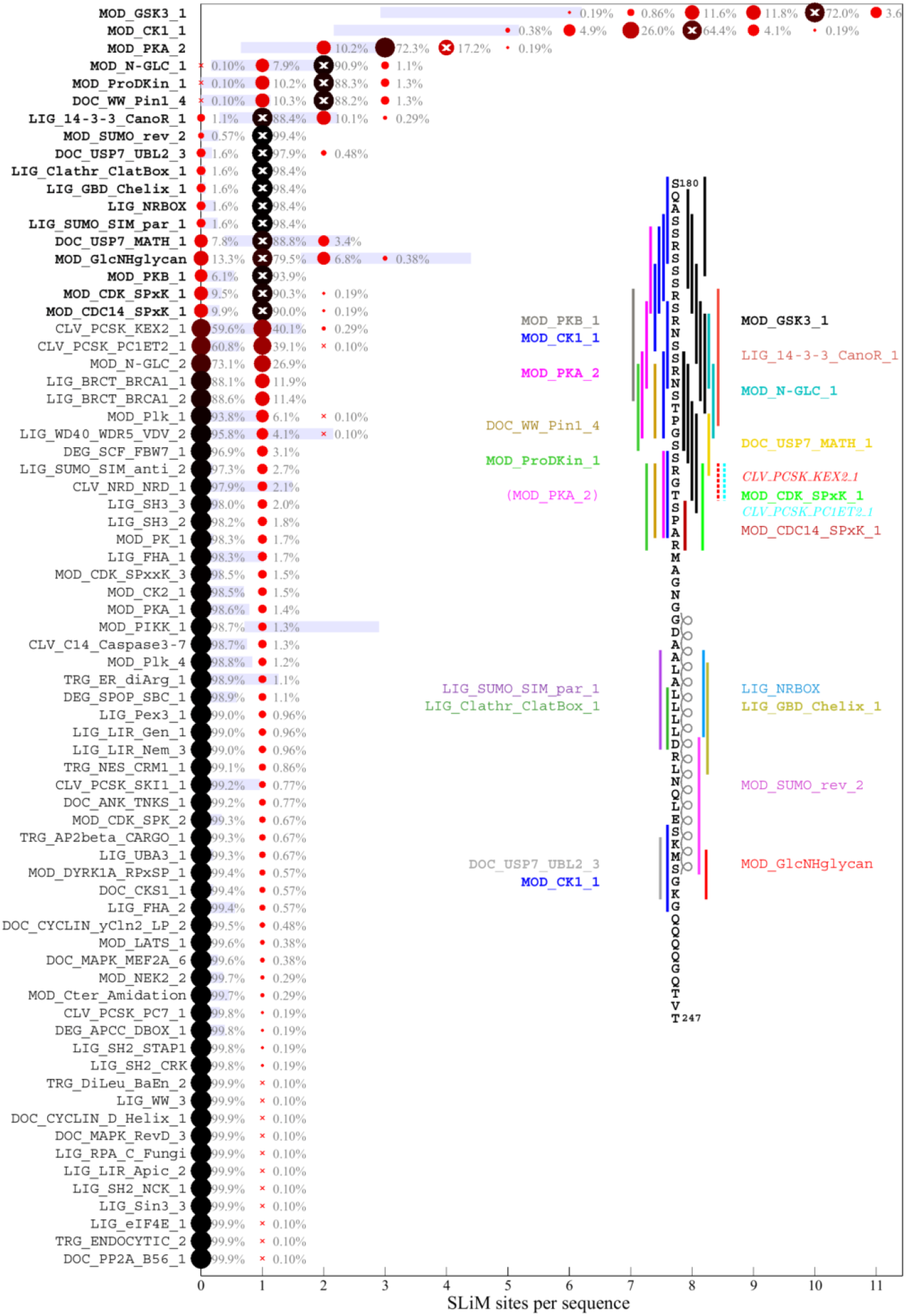
Predicted SLiM diversity in the linker IDR. Each row presents a histogram for the number of sites (abscissa) of a motif class in the ancestral reference sequence (bold) or in any of the 979 distinct mutant linker sequences. The frequency of the site numbers across the ensemble of sequences is indicated by symbol size and color, as well as the listed percentage value. The motif site number in the ancestral sequence is indicated by a white cross. The blue bars represent the mean ± standard deviation of the abundance of each motif in 10,000 randomly permutated reference sequences. The inset shows the location of the ancestral motifs as vertical bars and correspondingly colored motif name. New motifs emerging in the Omicron variant due to the defining R203K/G204R mutation are indicated as dotted lines and italicized motif name, and a disappearing motif is indicated in parenthesis.

**Figure 3.**
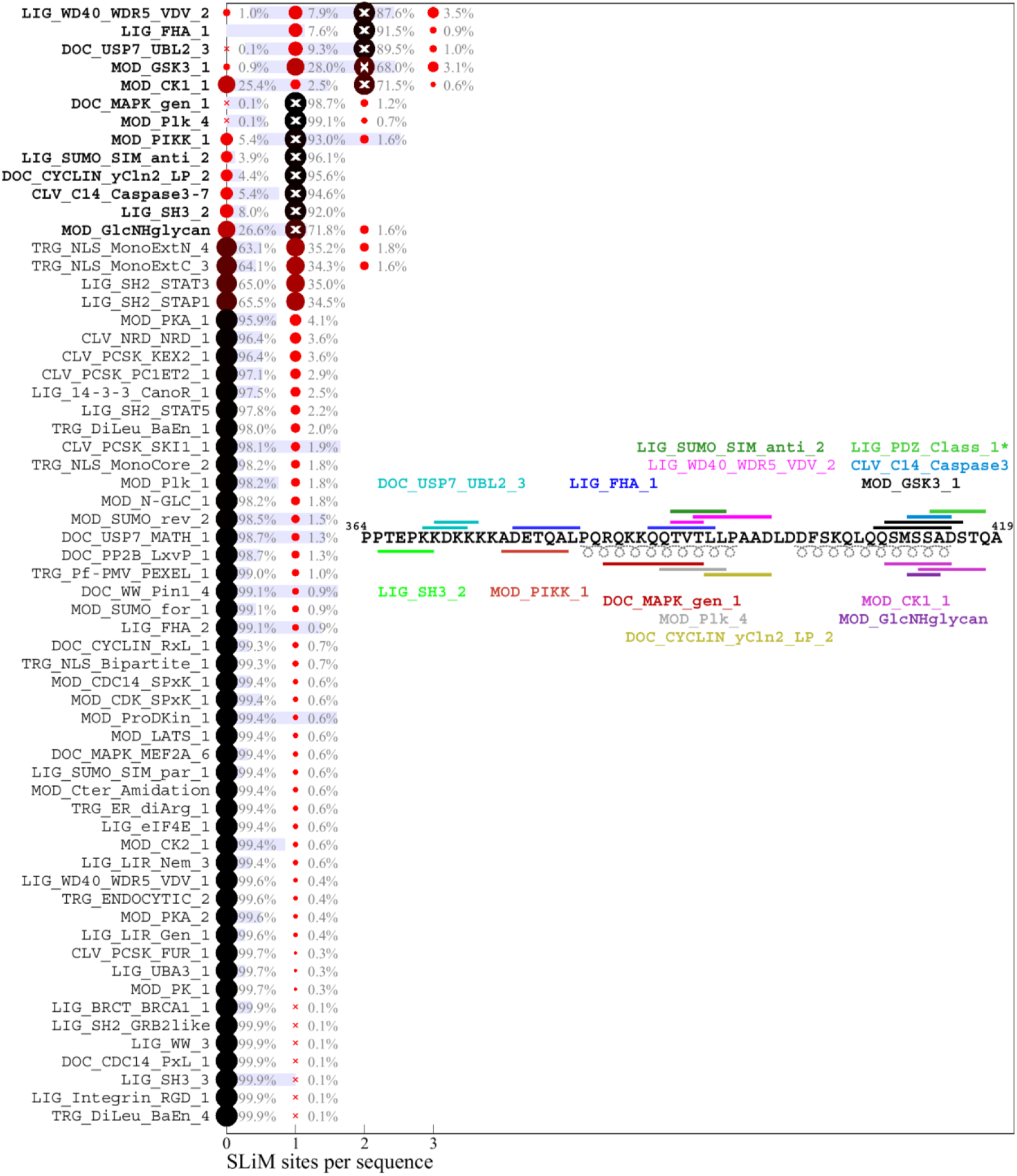
Predicted SLiM diversity in the C-arm IDR. Histograms for the number of sites of motif classes in the ancestral reference sequence (bold) or in any of the 556 distinct mutant C-arm sequences. Symbols and labels are as in Figure 2. The inset shows the location of the ancestral motifs as horizontal bars and correspondingly colored motif name. *LIG_PDZ_Class_1 is excluded from the distribution analysis (see **Methods**).

**Figure 4.**
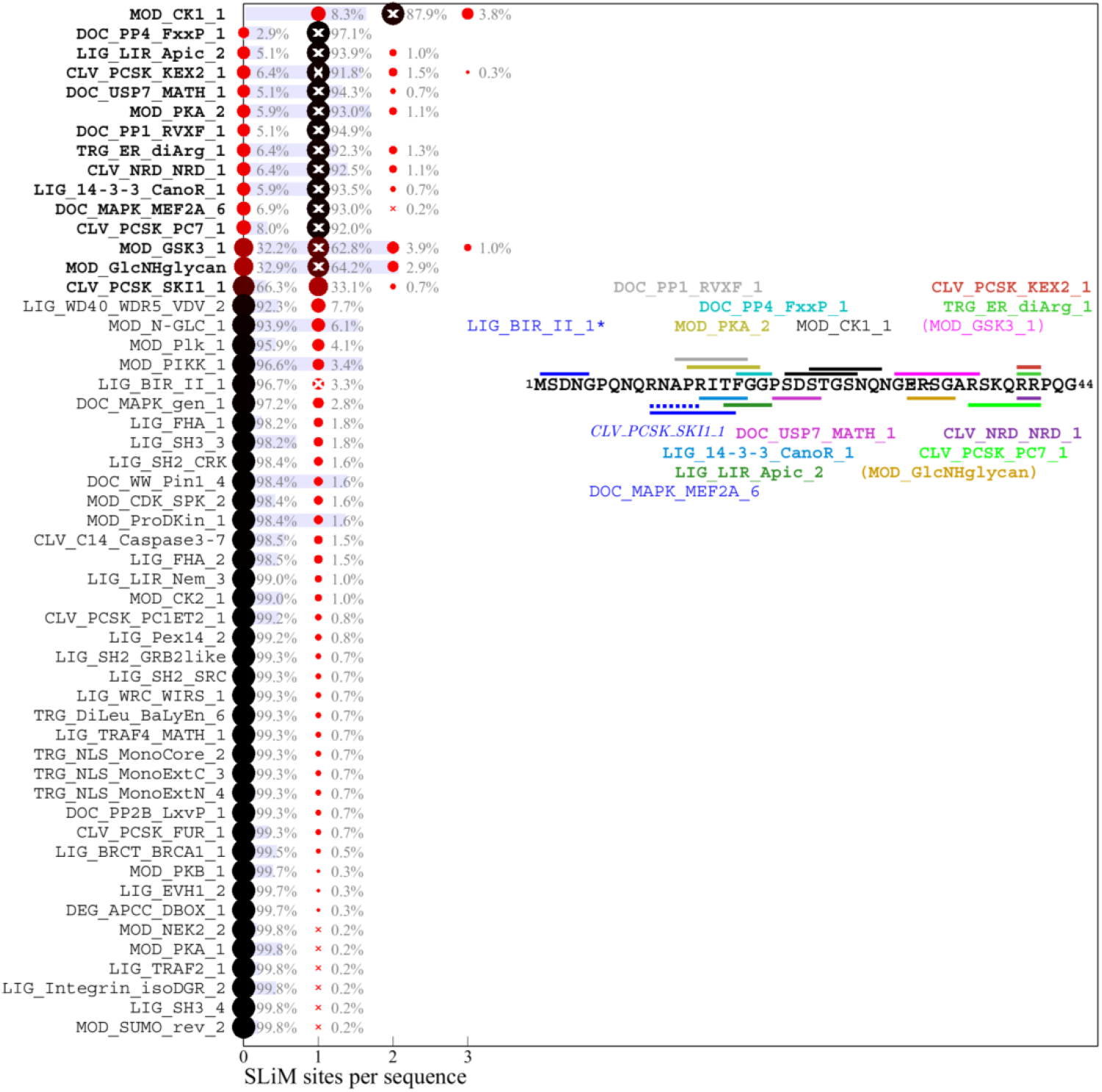
Predicted SLiM diversity in the N-arm IDR. Histograms for the number of sites of motif classes in the ancestral reference sequence (bold) or in any of the 512 distinct mutant N-arm sequences. Symbols and labels are as in Figure 2. The inset shows the location of the ancestral motifs as horizontal bars and correspondingly colored motif name. Omicron sequences have a defining substitution P13L and deletion in position 31-33. As a consequence, motifs in parenthesis do not occur in Omicron sequences, and motifs written in italics emerged. *LIG_BIR_II_1 is excluded from the distribution analysis (see **Methods**).

As a measure for the likelihood that some of these motifs may appear just stochastically we employ a strategy previously developed by Hagai and co-workers (Hagai et al., 2014): For each of the IDRs we generated a set of 10,000 randomly scrambled sequences with the same amino acid content, and for each motif, we determined the frequency of it occurring in the randomized set. The average number of sites with standard deviation is depicted as blue bars in **Figures 2-4**. As shown in the first three rows of **Figure 2**, there is a relatively high probability of generating multiple GSK3, CK1, and PKA phosphorylation sites in the linker IDR by chance, which may be expected given the amino acid composition particularly of the SR-rich region (**Figure 2**). However, their actual number in the ancestral sequence (white crosses) exceeds the statistical expectation, consistent with the important role of phosphorylation for intracellular N-protein (Carlson et al., 2020; Yaron et al., 2022). Other motifs of the ancestral sequence that have a high probability to occur by chance given the linker amino acid composition are sites for binding of 14-3-3 protein, USP7, and glycosaminoglycan attachment. Interestingly, a motif for interaction with yeast KEX2 protease has a high statistical chance to occur, and indeed is created through the defining R203K/G204R mutation of Alpha, Gamma, and Omicron variants. In the C-arm and N-arm IDRs, the majority of motifs displayed in the ancestral sequence are also likely to occur by chance given the respective IRD amino acid compositions. Unfortunately, the statistical analysis breaks down for motifs utilizing amino acids not part of the ancestral sequence, such as the defining 215C mutation of Delta variant linker, which creates a likely non-functional N-glycosylation motif.

### Distribution of motifs in the mutant spectrum of SARS-CoV-2 N-protein

The distribution of motifs across the observed sequences can be depicted as a histogram of the site multiplicity. Accordingly, for each motif in **Figures 2-4**, the color and size of the circles is scaled from large black to small red according to the frequency of sequences exhibiting different numbers of instances of that motif, as indicated numerically. A complete list of SLiMs and their frequencies can be found in the **Supplementary Information**. Strikingly, the major phosphorylation motifs in the linker (**Figure 2**) exhibit a great polydispersity in site numbers, for example, ranging from 6 to 11 with a mode of 10 for GSK3, from 5 to 10 with a mode of 8 for CK1, and from 2 to 5 with a mode of 3 for PKA sites. The sum of phosphorylation motifs ranges from 15 to 27. Even though a higher than statistically expected number of phosphorylation motifs is conserved across all sequences, it appears as if the detailed phosphorylation events are not critical for viable virus, as judged by the fact that individually none of the phosphorylation sites in the SR-rich region of the linker IDR are conserved in the mutational landscape (**Figure 1F**). Interestingly, most of the sequences have one less predicted PKA site than the ancestral sequence, which is caused by defining mutations R203K/G204R in Alpha, Gamma, and Omicron, and the R203M mutation in Delta variant; these mutations has have been shown experimentally to cause reduced phosphorylation and enhanced assembly functions and were hypothesized to reflect viral evolution (Syed et al., 2023).

A similar picture of prominent polydispersity emerges in the motif distributions of the C-arm and N-arm IDRs. Regarding the likelihood of motifs occurring by chance given the amino acid composition of the linker IDR, it is interesting to note that motifs with high statistical chance are indeed found in greater numbers in sequence subsets, which may be discerned for the glycosaminoglycan attachment, 14-3-3 binding, and USP7 binding motifs in the linker, as well as the GSK3 binding and glycosaminoglycan attachment sites in the N-arm.

A striking aspect of the motif content in the mutant spectrum is that nearly all motifs appear dispensable, which is indicated in the distinct sequence populations with zero sites in **Figures 2-4**. The only exceptions are the phosphorylation motifs of the linker and N-arm, the FHA binding motif in the C-arm IDR, and probably the G3BP1 binding motif in the N-arm not included in the ELM resource, as judged by the strong conservation of F17 in the mutational landscape. All other motifs are absent in sometimes sizeable fractions of mutant virus sequences, ostensibly suggesting that they may not describe real host protein interactions, or that these are not an essential part of the virus-host interface (see **Discussion**). For example, 1.1% of all linker sequences lack the 14-3-3 site, and 9.9% lack the CDC14 phosphatase dephosphorylation site, 25.4% of C-arm sequences lack both CK1 phosphorylation sites, and 32.2% of N-arm sequences lack the GSK3 site. Similarly, the defining Δ31-33 mutation in Omicron sequences destroys a likely non-functional glycosaminoglycan motif.

Conversely, many motifs that do not exist in the ancestral sequence are formed *ex nihilo* in subsets of sequences due to their particular constellation of mutations. In addition to several motifs arising from defining mutations of variants of concern (such as the yeast KEX2 protease site mentioned above), a large number of motifs occur in only a small fraction of sequences. However, since only sequences occuring in at least 10 different genomes were included in the analysis, a frequency of ≈1% of all linker sequences translates to on the order of 100 instances in the genome database.While many of those are plausible host protein interactions, others are less likely and may be random matches with regular expressions.

### Estimated accessible sequence and motif space of N-protein IDRs

The total number of motif classes displayed in the database sequences is 72 in the linker, 62 in the C-arm, and 53 in the N-arm IDR, out of a total of 327 motif classes currently contained in the ELM database. This raises the question of how efficiently random mutations can create new motifs, and what fraction of the total currently known motif space is accessible to N-protein. Since the viable amino acid landscape (**Figure 1F**) has been exhaustively explored during the pandemic so far, it is possible to make a back-of-the-envelope estimate of the theoretical maximal size of the associated sequence space by permutation through viable amino acid mutations at each position. For the N-arm (1:44) – the smallest of the N-protein IDRs – allowing for 3 mutations per sequence, which is close to the average of ≈2.8 observed in the existing database, there are ≈1.8×10^6^ different mutant sequence permutations. Although likely not all will be viable due to epistatic effects, this upper limit is more than three orders of magnitude larger than the 512 distinct N-arm sequences observed so far in the genomic database, and far exceeds our capacity for computational determination of the associated motif space.

Nonetheless, limited approximate sequence spaces can be searched when focusing on only the most frequently encountered amino acid mutations. For example, considering only the mutations observed in > 1,000 instances (i.e., in 0.02% of genomes; depicted in bold red in **Figure 1F**), the associated sequence space with three mutations consists of 10,500 sequences, which is searchable and describes 26 motif classes from the ELM database, nearly twice the number of different motifs in the ancestral N-arm sequence. Lowering the mutation frequency threshold leads to a rapidly growing sequence space and associated motif space (**Figure 5**). For example, at a mutation threshold of > 100 instances (mutation frequence of 0.002% in all genomes) 231,519 possible sequences with three mutations cover a range of 85 motif classes, or 26% of all in the ELM database. Consideration of more rare mutations occurring in the viable amino acid landscape further increases the theoretical sequence space and thereby the accessible motif space.

**Figure 5.**
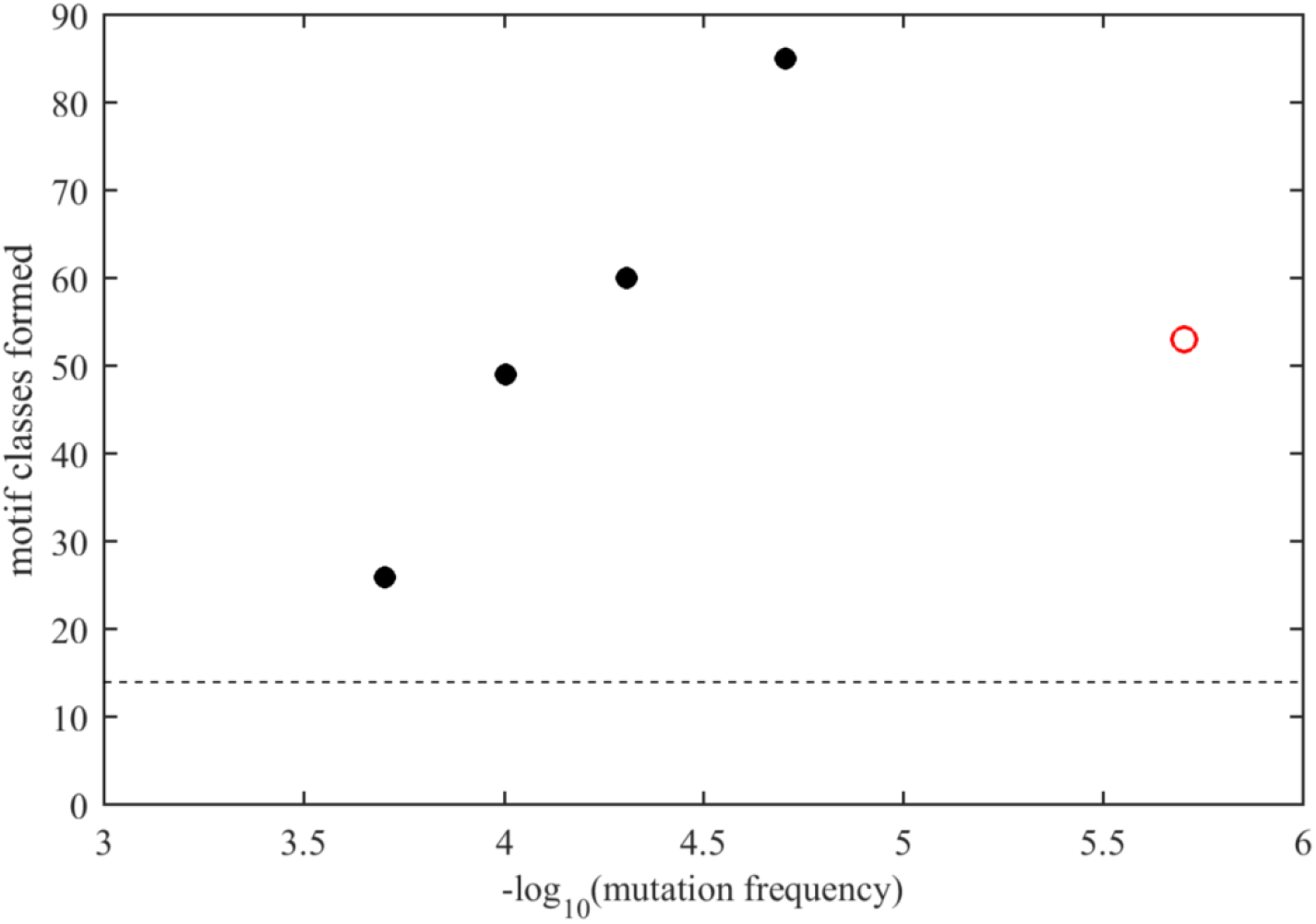
Estimated accessible motif space of N-protein N-arm IDR. Number of motif classes identified from the ELM database presented in the N-arm IDR examining a theoretical sequence space formed by three mutations per sequence permutated from the amino acid landscape (Figure 1F), considering only amino acid replacements with minimum frequencies indicated in the abscissa (number of observed mutation instances relative to a total of 5.06 million sequences). Black circles are completely evaluated theoretical sequence spaces, and the red circle is based on 512 distinct N-arm sequences contained in the GISAID database, comprising ≈0.03% of the theoretical sequence space. The dashed horizontal line is the number of motif classes displayed in the ancestral N-arm sequence.

## DISCUSSION

The virus-host interface is crucial for viral survival and a promising area for the development of antiviral therapeutics. Significant recent advances were driven by increased understanding of the evolutionary role of IDRs and the recognition of SLiMs as ubiquitous interaction modules that can be hijacked through viral mimicry. Unavoidably, both large-scale experimental and bioinformatics studies contributing to this picture were limited to consensus sequences of viral species, lacking opportunity to account for the quasispecies nature of RNA viruses, and thus leaving the salient feature of sequence diversity previously unexplored with regard to the role of SLiMs in the virus-host interface. However, an important recent development is the assembly of a vast SARS-CoV-2 genomic repository at GISAID (Elbe and Buckland-Merrett, 2017), which exceeds the number of available influenza sequences by more than one order of magnitude. This allows for the first time the exhaustive characterization of the amino acid mutation landscape (Bloom et al., 2023; Saldivar-Espinoza et al., 2023; Zhao et al., 2022), and is exploited here to power an analysis of the viral sequence space. This illuminates the highly dynamic motif space of viral IDRs, providing new insights in the unexpected efficiency of *ex nihilo* motif creation, the extent of viral motif mimicry, and the potential size of the virus-host interface.

We have started the present work with the assembly of the amino acid mutational landscapes from the host population-wide ensemble of consensus sequences. In a first approximation, we may consider the amino acid landscape as a reflection of all mutations consistent with vital biophysical, functional constraints of the viral protein, from which random sequence samples arise with certain mutation frequencies. Grouping all SARS-CoV-2 N-protein sequences in three major waves from different periods of the pandemic, groups essentially representing independent repeats of deep mutational scans, produces nearly identical mutational landscapes and local mutation frequencies. This suggests that the basic biophysical properties of N-protein overall have not significantly changed, as in an evolutionary stable steady-state. In the first approximation, this view justifies considering the derived motif space to be similarly in steady-state, and to represent an intrinsic property of N-protein.

This is notwithstanding fitness modulations from localized N-protein such as 203K/204R (Syed et al., 2023) and 215C (Zhao et al., 2022) that seem secondary to evolution of the immunodominant spike protein. Interestingly, in both Delta and Omicron waves the defining mutations lead to the destruction of one PKA motif that may impact the extent of linker phosphorylation and modulate the switch between intracellular and assembly functions (Syed et al., 2023). On the other hand, the observed variation in the content of kinase motifs is very large, and none of the potential phosphorylation sites other than S184 is strongly conserved in the mutational landscape, apparently without compromising virus viability. This points to a distributed phosphorylation threshold in the linker IDR rather than specific structural requirements (Zarin et al., 2021) for N-protein to be viable, which may be fine-tuned for fitness optimization.

The data for the available time-scale depict highly parallel random exploration of many motifs. On the level of single proteins, multiple overlapping repeat instances of motifs are displayed along the IDRs, a feature frequently encountered RNA viruses (Mihalič et al., 2023), which may lead to cooperativity and effective enhancement (Klein et al., 2003; Watson et al., 2022). Similarly, different overlapping motifs provide multi-functionality with little competition due to the large expression level with 10^8^ copies of N-protein in the infected cells (Tugaeva et al., 2021). Across the mutant spectrum, we observe highly effective motif creation, spanning an astonishingly wide range theoretically covering 20% or more of the known eukaryotic motif space even in the shortest N-protein IDR. Conversely, most of the motifs displayed in the ancestral sequence are destroyed in at least a subset of the viable mutant spectrum.

The origin of the sequence diversity in the consensus sequences considered here is rooted in the error-prone transcription and the intracellular quasispecies. However, it is unclear to what extent the observed SARS-CoV-2 sequence space reflects intracellular quasispecies and intrahost ‘meta-quasispecies’ (Domingo, 2019), perhaps constituting a ‘hyper-quasispecies’ as the ensemble of consensus sequences across the host-population, with the obvious exclusion of non-viable species. Even though the amino acid mutation landscape (**Figure 1F**) is sufficiently sampled to reflect biophysical features (Zhao et al., 2023), only a very small fraction of the implied theoretical sequence space has been sampled. Even less is known about the quasispecies, with deep sequencing capabilities probing intrahost minority sequences currently limited to a level of 0.1% (Martínez-González et al., 2022). However, several studies report that most of the mutations of intrahost minority species are independently reflected in the GISAID repository of consensus sequences (Martínez-González et al., 2022; Siqueira et al., 2021; Tonkin-Hill et al., 2021), demonstrating at least significant overlap. If the motif space observed in the present work is any indication of intracellular diversity, this would strongly further leverage the viral motif range. For example, it raises the possibility of cooperation between species (Vignuzzi et al., 2006), and one could envision minor species to interact with host proteins through motifs that the master sequence has abandoned. Thus, interaction motifs that disappear in a subset of consensus sequences in our study may nonetheless still be essential for viable virus. Also, the simultaneous presence of viral protein species with different motif sets attacking host processes at multiple entry points may exert synergistic effects on redundant interaction networks (Levy et al., 2010).

Whether the observed motif diversity extends intracellularly or not, the present work demonstrates that single sequence-based studies of viral protein/host interactions under neglect of viral diversity likely vastly underestimates the abundance of SLiM-based protein-protein interactions in the virus-host interface. On the other hand, many of the displayed motifs may not be functional host protein interactions, due to incorrect sequence context, localization, or even species specificity. This does not diminish the role of the dynamics of random motif generation, which may serve as a fertile basis to search for beneficial interactions for further optimization and host adaptation. Non-functional interactions as a consequence of ‘evolutionary noise’ may have little fitness penalty and should be expected to occur, as pointed out by Levy and colleagues (Levy et al., 2009), and this should hold true in particular for viral quasispecies.

## METHODS

### Mutational landscape

Mutation data were based on consensus sequences of SARS-CoV-2 isolates submitted to the GISAID, and downloaded on January 20, 2023 as database files metadata.tsv and nextclade.tsv preprocessed by Nextstrain (Hadfield et al., 2018). These contained ≈7.23 million genomes, of which only 5.06 million high quality sequences based on multiple criteria evaluated in the Nextstrain workflow were included here. As described previously (Zhao et al., 2022), for inspection of the mutational landscape a threshold of 10 observations of any mutation was used to filter adventitious sequencing errors. 746 sequences exhibiting insertions in the N protein were omitted. The Wuhan-Hu-1 isolate (GenBank QHD43423) (Wu et al., 2020) was used as the ancestral reference. Alignment of SARS and related betacoronavirus sequences was carried out with COBALT at NLM (Papadopoulos and Agarwala, 2007). SARS-CoV-2 sequences were grouped in sets of Omicron variants, Delta variants (Nextstrain 21J and later Delta clades), and sequences preceding 21J including Alpha, Beta, and other variant as well as ancestral sequences (termed pre-Delta). Processing, plotting, and analysis of the sequence data was performed with MATLAB (Mathworks, Natick, MA).

### Prediction of N-protein disorder and visualization

The N-protein structure was predicted using ColabFold (Mirdita et al., 2022). Since no confidence is achieved for residue angles in disordered regions, these were artificially stretched closer to 180° for better visualization. The resulting structure was plotted using ChimeraX (Pettersen et al., 2021) and colored according to the predicted IUPred3 score (Erdos et al., 2021).

### Analysis of SLiMs

SLiMs pattern recognition was carried out on distinct amino acid sequences with mutations in the disordered regions of interest, i.e., the N-arm (1-44), linker (180-247) and C-arm (364-419). To this end, all 5.06 million sequences were classified according to their N-protein IDR amino acid sequences. 512 distinct classes of N-arm sequences, 979 distinct linker sequences, and 556 C-arm sequences were identified that occurred in more than a threshold of 10 genomes.

In a preprocessing step, deletions were removed, sequence regions of interest were extended by 10 aa and, for efficiency to reduce server traffic, concatenated with AAAAA spacers to create composite sequences of length ≈1,000 aa prior to submission to the Eukaryotic Linear Motif Resource server (http://elm.eu.org) (Kumar et al., 2022). This concatenation process excludes from the statistics all N- and C-terminal-specific SLiMs, such as LIG_BIR_II_1 at the N-terminus of the N-arm and LIG_PDZ_Class_1 at the C-terminus of the C-arm, but avoids creating artificial termini at the limits of the IDR sequences of interest. System CURL commands were issued from a MATLAB script to send and receive data, carry out communication error control, and to parse the returned text to extract motif data. Motif content was mapped back and cropped onto the original sequence framework of interest, and motif name, multiplicity, and starting and ending positions tabulated for statistical analysis. Motifs with alternate regular expressions that completely overlapped were counted as a single instance. The frequency distribution of motif multiplicities across the different IDR sequence classes were derived from the table of motif multiplicity in each sequence.

To assess the probability of any SLiM occurring by chance, motif searches on 10,000 random sequences matching the amino acid content of the ancestral sequence of disordered linker and arms, respectively, were carried out. This was accomplished by performing random permutations reassigning amino acids to random positions within each IDR. The resulting sequences were analyzed for their motif content using the same computational pipeline as described for the sequence data above.

The analysis of hypothetical sequence space was carried out for the N-arm on the basis of the ancestral Wuhan-Hu-1 sequence, and the mutational landscape of observed amino acid mutations. In a first step, a set of possible mutations was created given a threshold of instances (or mutation frequency) for mutations to be considered. This creates an amino acid mutation table *a_pm_* of possible replacements of the ancestral residue by mutation *m* at position *p,* with 0 ≤ *m* ≤ *M_p_*, where *M_p_* is the total number of mutations above the threshold frequency *f* at that position *p*. In a second step, the set of possible sequences with *3* mutations was created in MATLAB by permutation through all combinations (*a_ix_*, *a_jy_, a_kz_*) with *i* < *j* < *k* with 1 ≤ *x* ≤ *M_i_*, 1 ≤ *y* ≤ *M_j_*, and 1 ≤ *z* ≤ *M_k_*, avoiding redundant symmetric permutations. The resulting amino acids at positions *i*, *j*, and *k* replaced the amino acids in the ancestral sequence. To determine the motif space associated with the accessible sequence space, each of the resulting sequences was subjected to the same motif analysis pipeline described above. This analysis was repeated for different threshold frequencies, which creates an extended set of considered amino acid mutations. For efficiency in the analysis of next lower threshold frequencies, only the new sequences not already contained in the previous set of higher threshold mutations were subjected to motif analysis; and results were merged with those already obtained at the higher threshold.

## ACKNOWLEDGMENTS

We thank the GISAID, Nextstrain, and ELM Resource teams and its contributors for maintaining these tools and databases. This work was supported by the Intramural Research Program of the National Institute of Biomedical Imaging and Bioengineering (ZIA EB000095-02), National Institutes of Health. This work utilized the computational resources of the National Institutes of Health HPC Biowulf cluster for sequence analyses.

