## Supplemental Figure 1 for "Diversity of Short Linear Interaction Motifs in SARS-CoV-2 Nucleocapsid Protein"

### SUPPLEMENARY INFORMATION

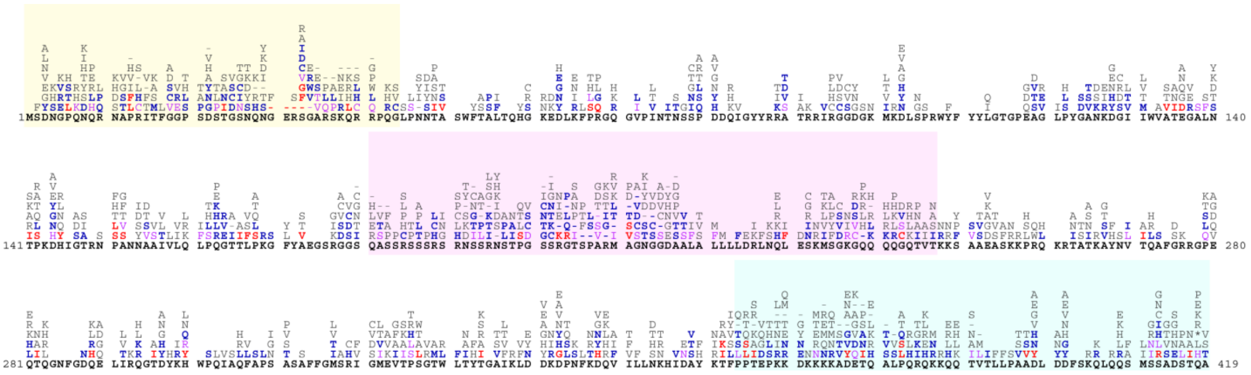

**Figure S1. Mutational landscape of SARS-CoV-2 Omicron N-protein**

As in **Figure 1**, shown in bold black letters is the ancestral amino acid sequence (Wuhan-Hu-1), and on top are mutations found in 5.05 million sequences from the GISAID repository, downloaded from Nextstrain in January 2023, and classified as one of the Omicron clades. The mutations are ordered and colored by number of genomes found to contain that specific mutation: 10 to 100 (gray), 100 to 500 (blue), 500 to 1,000 (purple), and > 1,000 (red). The intrinsically disordered regions are highlighted: N-arm in yellow, the linker in magenta, and C-arm in cyan.
